# Organ scale Three-dimensional atlas of developing pelvic innervation for the mouse lower urinary tract

**DOI:** 10.64898/2026.11.03.764023

**Authors:** Thomas M. Mehaffey, Reese Smith, Alan M. Watson, E. Michelle Southard-Smith

## Abstract

Pelvic autonomic neurons are responsible for coordinating pelvic organ function, including storage and emptying of the urinary bladder. These neurons arise from sacral neural crest-derived progenitors which migrate to the developing urogenital sinus and undergo differentiation alongside pelvic organs, ultimately organizing into paired pelvic ganglia. To circumvent the complex morphological changes that occur during genitourinary organ development, we have optimized light sheet fluorescence microscopy imaging for three-dimensional mapping of the development and spatial distribution of pelvic autonomic neurons. To track neural crest progenitor migration, neurogenesis, and formation of mature pelvic ganglia morphology in mice, we imaged intact lower urinary tract tissue from mid-gestational to post-weaning stages at single-cell resolution. The resulting atlas identifies the precise timing and initial routes of neural crest progenitor migration, defines timing for pelvic ganglion neurogenesis, and reveals previously unrecognized neuronal cell bodies outside of the centralized pelvic ganglia structures. The analysis further highlights morphological changes to pelvic ganglion structure that occur in parallel with genitourinary maturation, including the divergence of the major pelvic ganglia and bladder-proximal accessory ganglia. This work acts as the first wholemount imaging atlas of pelvic innervation in the mouse spanning fetal and postnatal timepoints, and thus we present an informative tool for future studies of developmental neurourology and neurogenic bladder dysfunction.

## INTRODUCTION

Autonomic and somatosensory innervation of the pelvis is necessary for the coordination of several processes, including urinary bladder micturition (emptying) and continence (storage), reproductive function, and defecation. The neurons which comprise pelvic-innervating structures arise from sacral neural crest-derived progenitor cells during development, migrating towards the developing lower urinary tract (LUT) and differentiating into diverse populations, including neurons with sympathetic and parasympathetic character.^1,2^ Despite shared neural crest origin of pelvic innervation, the spatial organization of differentiated neurons in the developing pelvic region is highly variable across species.

In humans, pelvic innervation is organized in a dispersed network of small ganglia positioned around the genitourinary organs known as the hypogastric plexus. In rodents, these neuronal structures are consolidated into larger paired pelvic ganglia which are positioned dorso-laterally on either side of the urethra flanking the bladder. The mature structure of the rodent pelvic ganglia includes multiple distinct neuronal clusters, with the largest population being the major pelvic ganglia (MPG), which are dorsolateral to the bladder neck and receive spinal input from the pelvic and hypogastric nerves (Figure 1). There are multiple postganglionic extensions from the major pelvic ganglia, including the cavernous nerves (also known as penile nerves in male and clitoral nerves in female) which extend posteriorly down the urethra (Figure 1). The rectal nerves also extend from the MPG dorsally towards the rectum, and a network of nerve fibers known as accessory nerves extend ventrally from the MPG towards the urinary bladder neck (Figure 1). Accessory nerves connect the MPG with smaller bladder-proximal clusters of neurons known as the accessory ganglia. Early molecular analysis of murine pelvic innervation classified postnatal neuron subtypes and early transcription factor expression in the MPG.^3,4^ However, because these studies most often dissect the MPG away from adjacent organs, the field is lacking a full spatial perspective of pelvic ganglia organization. Anatomical mapping of ganglia locations and histological estimates of neuronal size have been determined for the rat pelvic ganglia.^5^ However, these studies lack the three-dimensionality needed to gain full perspective of the structure of the pelvic ganglia. Analyses of rodent pelvic innervation have primarily focused on the male MPG, despite known sexual dimorphisms in the structure (Figure 1).^6,7,8^

**Figure 1.**
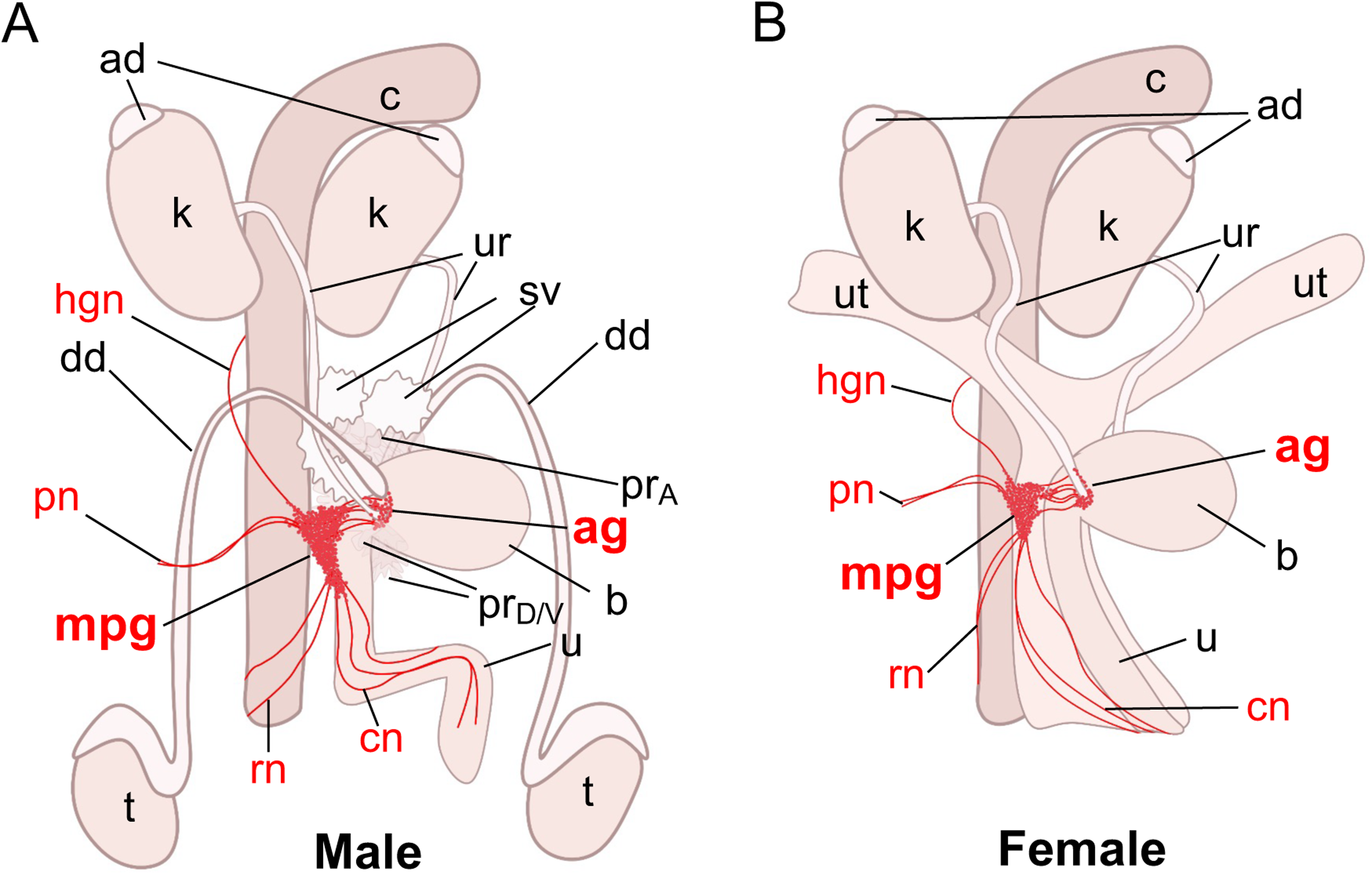
Anatomical location and morphology of the pelvic ganglia relative to LUT organs in the mouse. Schematized representation of (A) male and (B) female murine urinary tracts from a lateral view, with innervating structures represented in red. Abbreviations: **mpg,** major pelvic ganglia; **ag,** accessory ganglia; **b,** bladder; **u,** urethra; **k,** kidney; **ad,** adrenal gland; **ur,** ureter; **c,** colon; **sv,** seminal vesicle; **dd,** ductus deferens; **t,** testis; **pr_A_,** anterior prostate; **pr_D_,** dorsal prostate; **pr_V_,** ventral prostate; **ut,** uterine horn; **hgn,** hypogastric nerve; **pn,** pelvic nerve; **rn,** rectal nerves; **cn,** cavernous nerves.

Studies of pelvic innervation in the mouse largely focus on the anatomical organization and neuronal cell populations of the mature MPG. Relative to formation of other autonomic ganglia in the heart, lung, and sympathoadrenal axis, pelvic ganglia development has been relatively understudied. Historically, sacral neural crest-derived populations have been overlooked compared to other axial subpopulations of the neural crest, and the spatiotemporal organization of pelvic ganglia formation across developmental time has not been extensively reported. Multiple reports that included aspects of sacral neural crest migration did so in the context of analyzing contributions of this progenitor population to the enteric nervous system (ENS).^9,10,11,12^ Previous work from our group identified key timepoints and approximate routes for neural crest-derived progenitor cell migration and neuronal differentiation in the pelvic ganglia by tracking *Sox10*+ neural crest-derived progenitors.^3^ This analysis defined the migratory routes of sacral neural crest-derived progenitors, which give rise to the pelvic ganglia, at discrete timepoints during fetal mouse organogenesis; however high-resolution spatial imaging from the earliest stages when pelvic sacral progenitors initially aggregate was not assessed. Initial phases of sacral neural crest migration and formation of early neuronal aggregates of the pelvic ganglia are known to occur, at least in part, independently of preganglionic nerve input.^13^ While some aspects of these initial studies were performed in wholemount, the analyses lacked three-dimensional resolution and focused on lateral viewing perspectives. As a result, a comprehensive spatial perspective during pelvic ganglia formation in relation to the hindgut and organogenesis of the LUT is lacking.

In the present study, we sought to address these gaps by establishing a method for three-dimensional imaging of the mouse LUT with the pelvic ganglia positioned *in situ* alongside LUT organs *in situ*. In recent years, wholemount tissue clearing and imaging has provided improved perspective for many tissues and organ systems, with notable method development and applications in the brain. By applying tissue clearing and light sheet fluorescence microscopy (LSFM) to wholemount LUT tissue from postnatal day 28 (P28) mice, we were able to visualize relevant three-dimensional structures while retaining *in vivo* spatial organization. Using genetic tools to visualize neural crest lineages, we have produced the first three-dimensional renderings of the mouse pelvic ganglia, including intact MPG and accessory ganglia with relevant postganglionic extensions in place. With the added perspective supplied by LSFM, we identify intricacies in neuronal localization within the pelvic ganglia structure which have not previously been appreciated in the field. Further, by applying our optimized methods for tissue clearing and LSFM to the developing LUTs in fetal mice, we have produced an atlas of developing pelvic innervation that spans initial migration of sacral neural crest to mature pelvic ganglia morphology. Data presented here will act as a framework for the field of developmental neurourology and will offer context for future studies of the molecular underpinnings controlling neural crest progenitor migration, neuronal differentiation, and pelvic ganglia function.

## RESULTS

### 1. LSFM captures in vivo organization of the mature mouse pelvic ganglia

Given the complexity of mammalian pelvic innervation, and the relative paucity of studies that spatially map pelvic ganglia development, we sought to establish methods for tissue clearing and wholemount LSFM imaging of the mouse LUT. For protocol optimization, LUTs were collected from *Sox10-cre; R26R-tdTomato* mice at P28, providing fluorescent visualization of neural crest-derived cell populations, including the neurons and glia of the pelvic ganglia and associated nerve fibers. To maintain spatial integrity of the pelvic ganglia and adjacent innervated organs, and to avoid tissue shrinkage associated with many solvent-based tissue clearing approaches, we applied the aqueous CUBIC (Clear, Unobstructed Brain/Body Imaging Cocktails) method.^14^ This method consists of a two-part delipidation and refractive index matching approach to achieve optical clarity. Despite tissue swelling, which is a well-documented outcome of the lipid removal step,^14^ we observed that the CUBIC method caused less tissue distortion in the LUT than other clearing approaches, thus preserving the relative *in vivo* placement of pelvic ganglia neurons faithfully alongside target structures. We found that CUBIC clearing, in tandem with fluorescence-guided dissection to remove extraneous obstructing tissue, was sufficient to visualize the pelvic ganglia (Figure 2).

**Figure 2.**
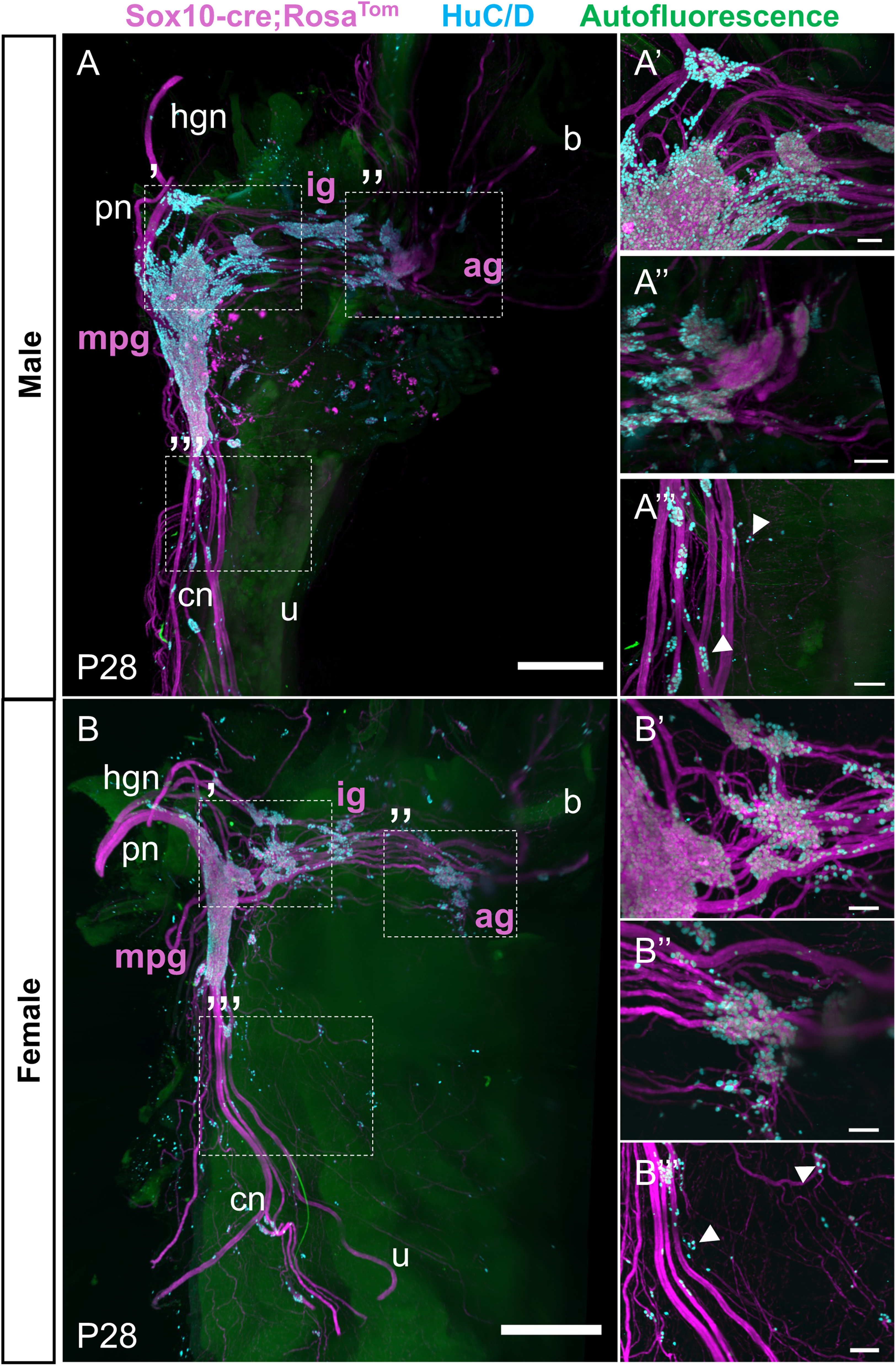
LSFM captures *in situ* organization of the mature mouse pelvic ganglia. LSFM images of wholemount lower urinary tracts from *Sox10-cre; R26R-tdTomato* (A) male and (B) female mice at P28 show the organization of pelvic innervation relative to target organs. Neural crest lineages, endogenously marked by the *Sox10-cre; R26R-tdTomato* transgene, are shown in magenta. Immunostaining for the pan-neuronal marker HuC/D is shown in cyan. Autofluorescence is shown in green. Signal from *tdTomato*-expressing glandular urethra tissue has been masked out to improve pelvic ganglia visualization in panel (A). Inset panels highlight the complexity of neuronal organization within the pelvic ganglia, with several HuC/D-positive neurons present in the anterior aspect of the major pelvic ganglia (A’, B’), in the bladder-proximal accessory ganglia (A’’, B’’), and down the length of the cavernous nerves extending posteriorly (A’’’, B’’’, white arrowheads). Abbreviations: **mpg,** major pelvic ganglia; **ig,** intermediate ganglia; **ag,** accessory ganglia; **b,** bladder; **u,** urethra. Scale bars = 1000um (A, B), 200um (inset zooms).

Through CUBIC clearing and LSFM, we identify intricacies in neuronal distribution throughout the LUT that have been historically overlooked. To visualize individual neuronal somata present on *Sox10-cre; R26R-tdTomato* LUTs, we performed wholemount immunohistochemical staining for the pan-neuronal marker HuC/D. At the anterior aspect of the major pelvic ganglia, neuronal cell bodies are present singularly and in small clusters up the length of the hypogastric and pelvic nerves (Figure 2A’/B’). As reported previously in rat,^5^ the bladder-proximal accessory ganglia are closely associated with the ureter entry sites in both males and females, and with the ductus deferens entry site in males (Figure 2A’’/B’’). These accessory ganglia are larger and more complex than has been previously appreciated with a multilobular appearance that is particularly prominent in males. There are also sizable clusters of HuC/D+ neuronal somata in ‘intermediate ganglia’ at the midpoint along accessory fibers that connect the major pelvic ganglia and accessory ganglia (Figure 2A/B). In both males and females, there are multiple individual neuronal somata and small ganglia associated with the cavernous nerve which extends posteriorly down the urethra (Figure 2A’’’/B’’’). There are also many neuronal somata present on the ventrolateral surface of the urethra which, to our knowledge, have not previously been described in the literature.

We also observe several sexual dimorphisms in pelvic ganglia structure and position from our LSFM imaging of P28 LUTs. While studies of the female mouse pelvic ganglia are limited, there is a large body of literature describing sexual dimorphism in the rat pelvic ganglia.^7,8^ In agreement with these studies, we find that the female mouse major pelvic ganglia are smaller than those of the male mouse. The female accessory ganglia and intermediate ganglia are also smaller than in the male. In both male and female mice, the position of the major pelvic ganglia is dorsolateral to the urethra (Figure 2A/B). The female major pelvic ganglia are positioned more dorsally than in the male, aligned laterally with the cervix behind the urethra (Figure 2B). As a result, the cavernous nerve extends more ventrally in female mice, while the cavernous nerve extends more posteriorly in male mice. As we have described previously, *Sox10-cre; R26R-tdTomato* expression is observed in glandular tissue of male and female mice, and this expression is independent of the neural crest lineages.^15^ While this tdTomato signal is present prominently in glandular tissue down the entire length of the male urethra (Figure S1A), this signal is confined to the distal region of the female urethra (Not shown). In P28 male mice, we also observe bright aggregates of tdTomato-expressing cells in the distal regions of prostate lobes (Figure S1A’).

### 2. Bladder-proximal accessory ganglia are comprised of a subset population of spatially distinct ganglia

While accessory ganglia populations have been described in detail in rat pelvic innervation,^5^ the structure of the bladder-proximal accessory ganglia in mouse has not previously been reported. Given our observations of the size and complexity of the accessory ganglia present at the bladder neck in P28 mice, we investigated the spatial organization of these ganglia more closely. In our LSFM images of optically cleared LUTs from P28 *Sox10-cre; R26R-tdTomato* mice, we consistently observe spatial separation between clusters HuC/D-expressing neuronal somata at the bladder neck in accessory ganglia and more dorsally situated clusters of neuronal somata in intermediate ganglia (Figure 3A). To improve our visualization of only the accessory ganglia, we defined the exact boundaries of the bladder-proximal ganglia and masked out signal from HuC/D+ cell bodies outside of these boundaries using the *Imaris* image analysis software (Figure 3A’, A’’). Once only neuronal somata within the accessory ganglia population were visible, we were able to define discrete sub-ganglia populations present at the bladder neck. In agreement with observations in the rat,^5^ we consistently see two large populations of accessory ganglia in male LUTs at P28: one positioned laterally, and one positioned closer to the midline, dorsal to the bladder neck (Figure 3B). These largest accessory ganglia populations are also observed in female LUTs (Figure S2). We also observe multiple smaller ganglia located immediately adjacent to the larger accessory ganglia in the mouse (Figure 3B). All discrete lobules of the accessory ganglia are significantly smaller than the MPG (Figure 2). Given their proximity to the bladder neck, there is very close association between the accessory ganglia and the ureters in male and female mice, as well as with the ductus deferens in males (Figure 1).

**Figure 3.**
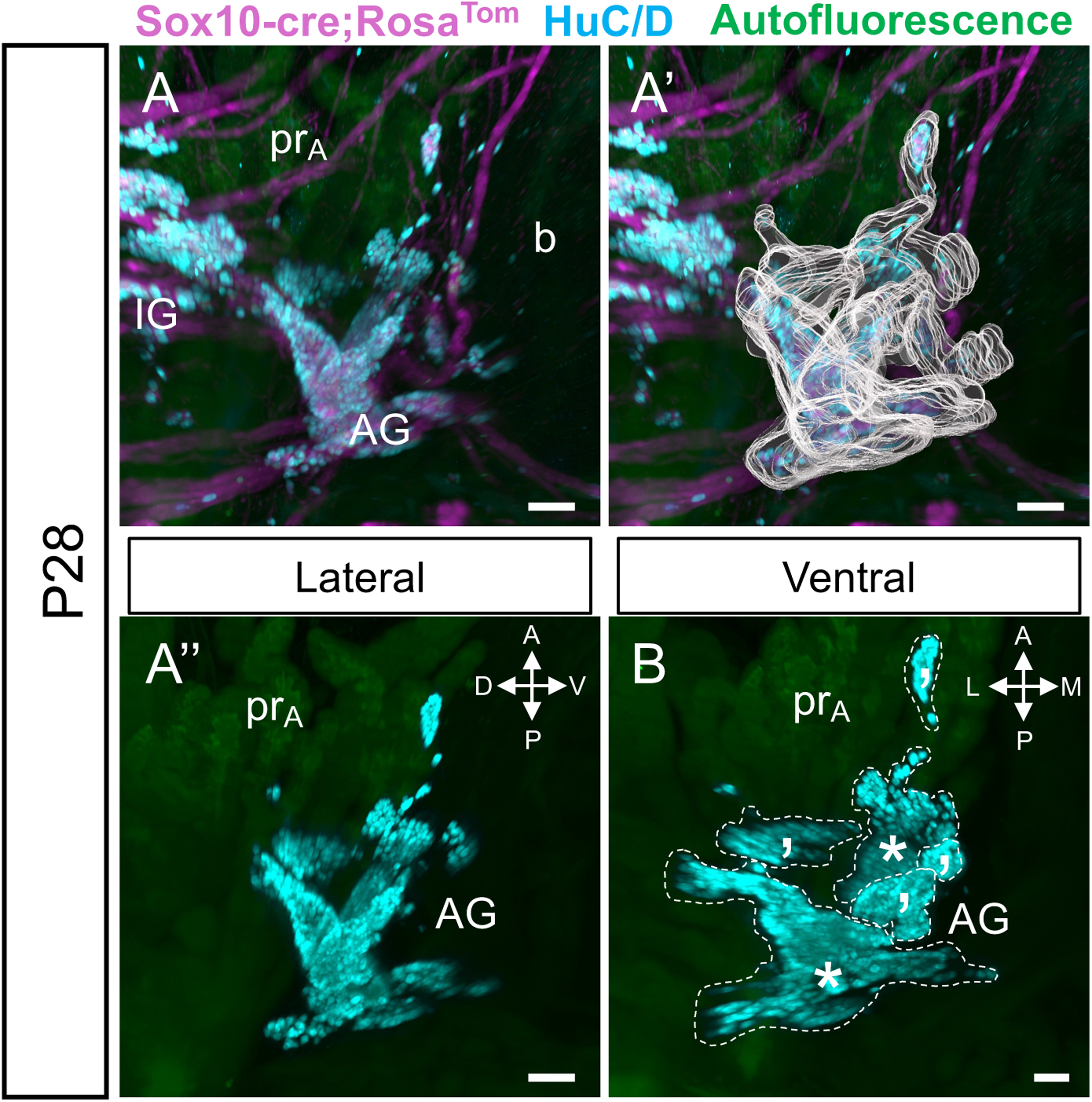
Bladder-proximal accessory ganglia are comprised of smaller, spatially distinct ganglia. Wholemount LSFM imaging of the lower urinary tract from a male *Sox10-cre; R26R-tdTomato* mouse at P28 reveals the complexity of the accessory pelvic ganglia on the right lateral surface of the bladder neck (A). Neural crest lineages (magenta) are shown relative to neuronal somata labeled by immunostaining for HuC/D (cyan). To improve visualization, spatial distribution of the accessory ganglia was defined according to HuC/D fluorescent signal (A’), and HuC/D positive neuronal somata outside of the defined region were masked out of the image to isolate the accessory ganglia (A’’). Three-dimensional observation of the accessory ganglia both from lateral (A’’) and ventral (B) perspectives shows discrete ganglia populations present at the bladder neck (dashed lines). Larger populations are marked with asterisks (*) and smaller populations are marked with apostrophes (’). Abbreviations: AG, accessory ganglia; IG, intermediate ganglia; b, bladder; pr_A_, Anterior prostate. Scale bars = 100um.

### 3. There is ongoing separation of spatially distinct pelvic ganglia subpopulations postnatally

We next sought to visualize how pelvic ganglia morphology changes as urinary and reproductive organs grow and mature postnatally. It has previously been reported that separation of early neurons into the distinct clusters of MPG and AG occurs between 17.5 days post coitum (dpc) and P10.^16^ However, the exact timing of when this process occurs has not been known. Male LUTs were collected from *Sox10-cre; R26R-tdTomato* animals at P2 and P10 for comparison to P28 (Fig 2A). Samples at these younger timepoints were processed through our optimized CUBIC clearing protocol, immuno-stained for HuC/D expression, and imaged via LSFM. At P2, just after birth, the cluster of neurons which comprises the major pelvic ganglia extends continuously from the lateral urethra to the bladder neck, with no obvious separation between the major pelvic ganglia and accessory ganglia (Figure 4A). We also see in P2 samples that there is a high prevalence of neuronal somata in small clusters on the ventral urethra. By contrast, at P10, the major pelvic ganglia and accessory ganglia are evident as distinct populations, and there is a tightly associated network of nerve fascicles connecting the separated ganglia (Figure 4B). However, at P10, there is no clear separation between the major pelvic ganglia and the intermediate ganglia that is observed at P28. Between P2 and P10, the shape of the major pelvic ganglia becomes more elongated, extending farther both posteriorly and ventrally. There is also a marked decrease in neuronal somata on the ventral urethra at these timepoints. After P10, remodeling of the pelvic ganglia continues, with separation between the major pelvic ganglia and intermediate ganglia occurring by P28 (Figure 4C). As the genitourinary organs grow between P10 and P28, nerve fascicles between the major pelvic ganglia and the accessory ganglia also become more dispersed. We observe the same continuum of change in pelvic ganglia morphology in female mice at postnatal timepoints (Figure S3).

**Figure 4.**
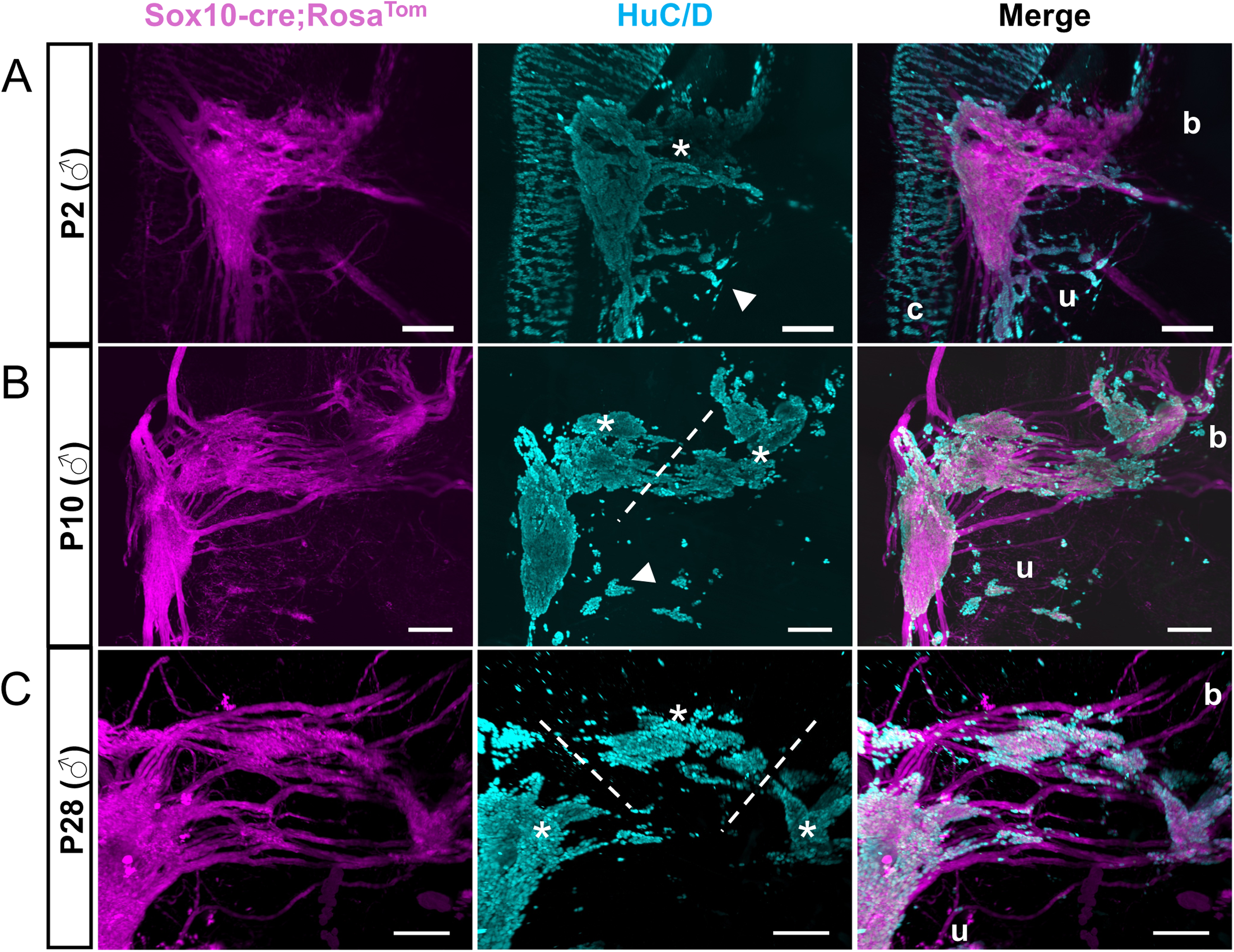
Separation of spatially distinct pelvic ganglia subpopulations occurs postnatally. Wholemount lower urinary tracts from *Sox10-cre; R26R-tdTomato* male mice imaged via LSFM at postnatal timepoints show a continuum of change in pelvic ganglia organization after birth. Neural crest lineages (magenta) are shown relative to neuronal somata labeled by immunostaining for HuC/D (cyan). At P2 (A), there is no discernable separation between the major pelvic ganglia and the accessory ganglia at the bladder neck (population marked by asterisk). By P10 (B), there is a clear distinction between the centralized major pelvic ganglia and the bladder-proximal accessory ganglia (populations marked by asterisks, separated by dotted line). Finally, at P28 (C), mature morphology of pelvic innervation is established, and a diffuse network of nerve fascicles connect separate populations of major pelvic ganglia, intermediate ganglia, accessory ganglia (populations marked by asterisks, separated by dotted lines). Abbreviations: **b,** bladder; **u,** urethra. Scale bars = 300um.

### 4. Neuronal populations expand to the ventral surface of the urethra during late gestation

Late gestation in the mouse is known to be a key period of growth in genitourinary organs, with rapid differentiation of anatomically distinct sex-specific organs.^17,18^ To examine the organization of pelvic ganglia neurons during this time, we collected LUT tissue from male and female *Sox10-cre; R26R-tdTomato* embryos at 15.5dpc and 18.5dpc. Previous studies have reported the presence of centralized clusters of neurons alongside the lateral surface of the urethra by 15.5dpc.^3,19^ However, these prior reports lacked the spatial resolution of wholemount LSFM and thus did not resolve the position of neurons relative to the urethra. By applying CUBIC clearing, immuno-staining for HuC/D, and LSFM, we observe that neuronal somata are present in high density to the lateral aspects of the urethra in the pelvic ganglia in male and female embryos at 15.5dpc (Figure 5A). Between 15.5 and 18.5dpc, there is a notable increase in small clusters of neuronal somata localized outside of the major pelvic ganglia. By 18.5dpc in male embryos, neuronal populations have expanded ventrally towards the bladder neck, posteriorly down the length of the urethra, and onto the ventral surface of the urethra (Figure 5B). The female embryo at 18.5dpc also exhibits an expansion in neuronal populations outside of the major pelvic ganglia, though there are fewer neuronal clusters located on the ventral urethra and more neuronal clusters positioned dorsally and posteriorly than in the male (Figure S4).

**Figure 5.**
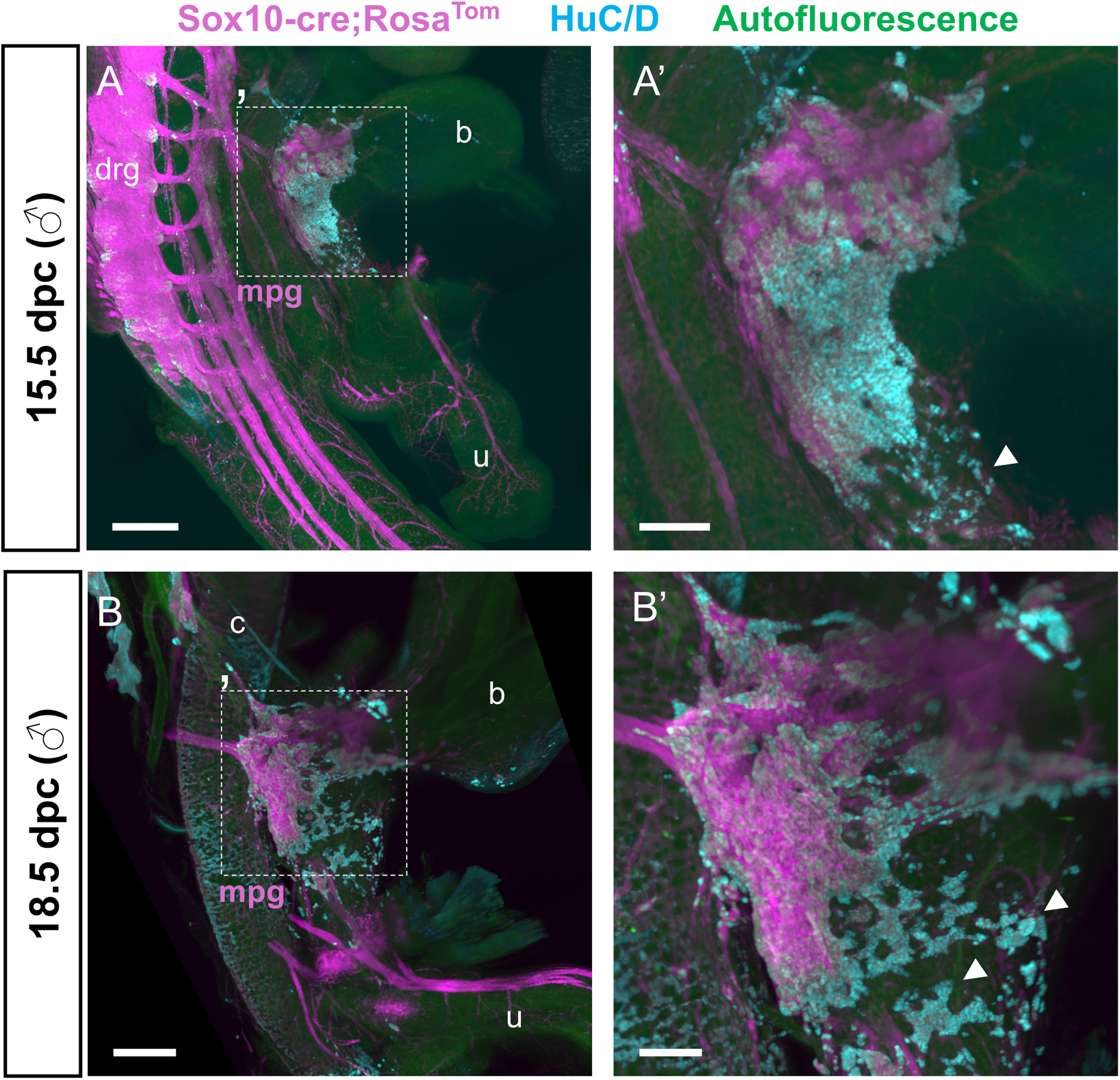
Neuronal populations expand to the ventral surface of the urethra during late gestation. Wholemount lower urinary tracts from *Sox10-cre; R26R-tdTomato* embryos at 15.5dpc (A) and 18.5dpc (B) show changes in neuronal cell body localization in the pelvic ganglia which occur at late embryonic timepoints. Neural crest lineages (magenta) are shown relative to neuronal somata labeled by immunostaining for HuC/D (cyan). At 15.5dpc, a centralized pelvic ganglia structure has formed, with only a few neuronal cell bodies present on the ventral urethra (A’, white arrowhead). By 18.5dpc, just before birth, pelvic ganglia neurons have expanded such that the ventral urethra is covered in neuronal cell bodies (B’, white arrowheads). Scale bars = 500um (A,B), 200um (inset zooms).

### 5. Neural crest-derived progenitors positioned at the genital tubercle initiate neurogenesis by 12.5dpc

Having established differences in distribution of neuronal somata for pelvic ganglia from late gestation onward, we sought to capture a 3-D perspective of the early steps in LUT innervation, when neurogenesis first occurs. Previous studies have shown that sacral neural crest-derived progenitor cells are present laterally on either side of the urogenital sinus at the base of the genital tubercle by 12.5dpc.^3,9,13^ Wiese and colleagues also reported the expression of early pan-neuronal markers (HuC/D, TuJ1) in these progenitors at 12.5dpc. In the present study, we applied tissue clearing, HuC/D immuno-staining, and LSFM to *Sox10-cre; R26R-tdTomato* embryos at 12.5dpc, as this method provides improved spatial perspective for the distribution of HuC/D-expressing cells in the developing genital tubercle. With our wholemount LSFM method, we observe clusters of neural crest-derived progenitors alongside the lateral sides of the genital tubercle by 12.5dpc (Figure 6). Notably, we find that HuC/D expression in the neural crest-derived progenitor population at 12.5dpc is confined to the distal-most region of this migratory population, lateral to the genital tubercle. (Figure 6A’’). There are also *tdTomato-*expressing cells that extend laterally past the initial aggregate of neuronal progenitors towards the distal base of the genital tubercle (Figure 6A’), whether these long streams consisted of cellular processes or thin streams of cells was not clear due to the cytoplasmic distribution of *tdTomato*. To examine this aspect further and visualize individual migratory neural crest-derived progenitors, we also collected embryos expressing the *Sox10-H2BVenus* transgene, that express the nuclear localized YFP reporter.^20^ For added validation of early neurogenesis in the pelvic ganglia anlagen, we immuno-stained tissue from *Sox10-H2BVenus* embryos at 12.5dpc for HuC/D, localized in neuronal somata, and beta tubulin class III, or TuJ1, localized in neuronal projections. As with the *Sox10-cre; R26R-tdTomato* model, we see the initiation of neurogenesis in NCPCs alongside the genital tubercle by 12.5dpc, marked by both HuC/D and TuJ1 expression (Figure 6C).

**Figure 6.**
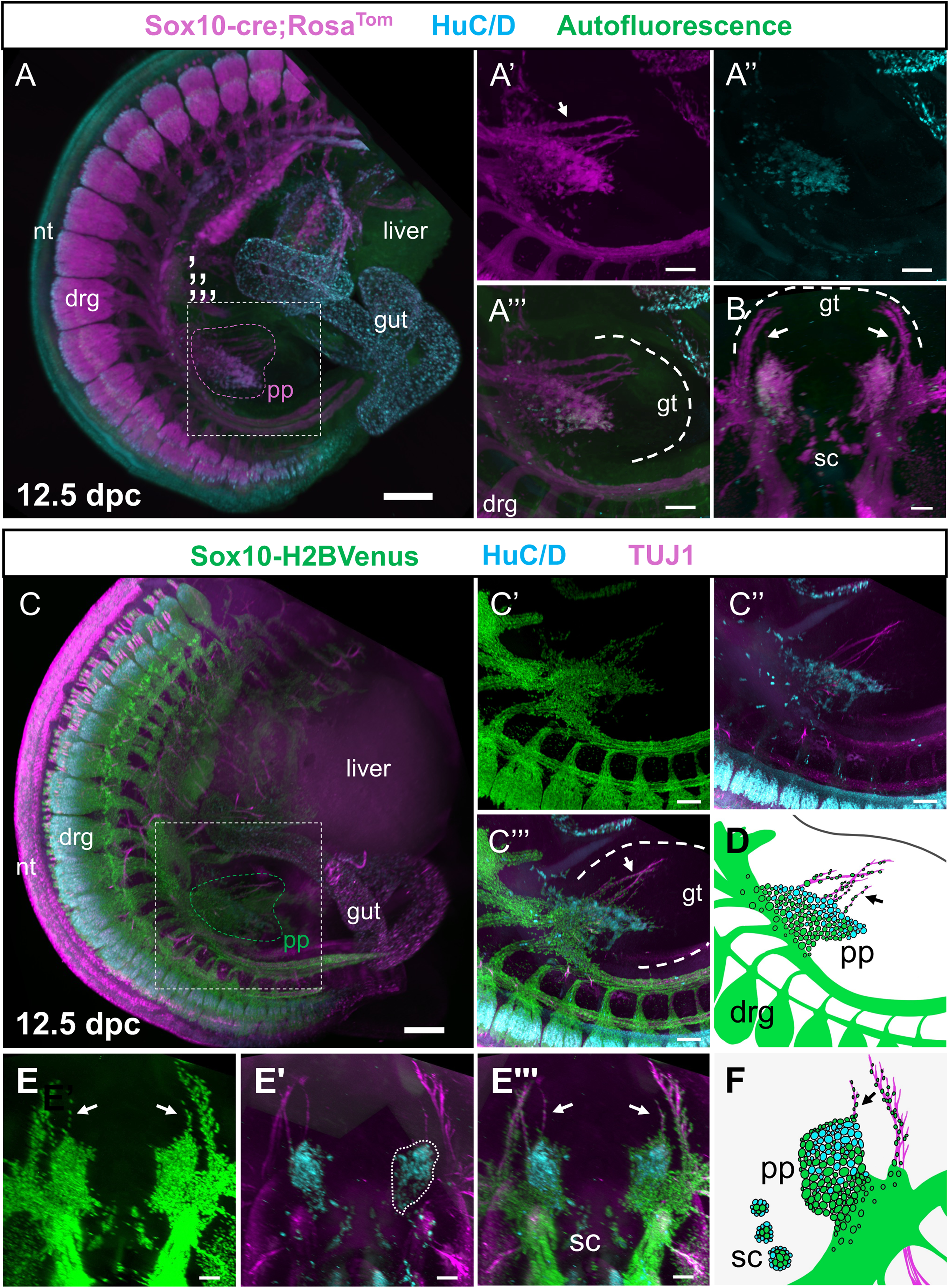
Neural crest-derived progenitors positioned at the genital tubercle initiate neurogenesis by 12.5dpc. Wholemount LSFM images of mouse embryos at 12.5dpc captures spatial distribution of neuronal progenitor cells alongside the developing lower urinary tract. In *Sox10-cre; R26R-tdTomato* embryos at 12.5dpc (A), neural crest lineages (magenta) are shown relative to neuronal somata labeled by immunostaining for HuC/D (cyan). Arrows point to neural fibers. At this timepoint, a population of neural crest derivatives is present on the lateral genital tubercle (A’), and cells within this population are expressing HuC/D (A’’). The transverse perspective of LSFM three-dimensional renderings (B) confirms that populations of neural-crest derived cells are present bilaterally on the genital tubercle. Imaging of *Sox10-H2BVenus* embryos at the same timepoint allows visualization of *Sox10*-expressing migratory neural crest progenitor cells, shown here in green (C). Immunostaining for HuC/D, shown in cyan, and TuJ1, shown in magenta, provides further perspective for the localization of neuronal somata and processes, respectively. As in the *Sox10-cre; R26R-tdTomato* model, neural crest progenitor cells are present at the lateral genital tubercle at this timepoint (C’), and cells in this population are expressing markers of neuronal lineage commitment (C’’). (D) Schematic emphasizes main elements in lateral views. The transverse perspective of the *Sox10-H2B-Venus* sacral region (E) also shows populations of neural crest progenitor cells on the left and right sides of the genital tubercle. (F) Schematic to emphasize elements in the transverse perspective. Abbreviations: nt, neural tube; drg, dorsal root ganglia; pg, pelvic ganglia progenitor cells; gt, genital tubercle. Scale bars = 500um (A,C), 200um (insets, B,E).

### 6. LSFM reveals precise spatiotemporal organization of pelvic ganglia formation and neurogenesis

To gain a better understanding of neural crest-derived progenitor migration and early changes that lead to formation of the neuronal aggregate alongside the genital tubercle, we collected embryos expressing the *Sox10-H2BVenus* transgene reporter at discrete timepoints between 11.0dpc and 12.5dpc based on somite staging of collected samples. Sacral neural crest migration is underway in the mouse by 10dpc,^3^ and as previously shown, neural crest progenitors are undergoing neurogenesis at the base of the genital tubercle by 12.5dpc. Despite prior efforts to map the migration pathways of the progenitors which give rise to the pelvic ganglia, the precise spatial routes and temporal control over migration are not well understood due to the complex morphological changes that occur during hindgut septation and formation of the urogenital sinus. By applying wholemount tissue clearing, immuno-staining for HuC/D and TUJ1, and LSFM, we have produced a map of sacral neural crest migration and neurogenesis with improved resolution and 3-D spatial perspective. By 11.0dpc, a small population of migratory NCPCs, marked by *H2BVenus* fluorescence, have moved bilaterally to the genital tubercle (Figure 7A/F). By 11.5dpc, these migratory populations have expanded, as progenitors begin to infiltrate the lateral surfaces of the growing genital tubercle (Figure 7B/G). Between 11.5 and 11.75dpc, these clusters of progenitors expand in cell numbers, and undergo a notable increase in the number of progenitors expressing HuC/D and TUJ1 (Figure 7C/H). By 12.25dpc, the cluster of progenitor cells has grown out towards the distal genital tubercle, with HuC/D positive cells positioned at the distal region of the progenitor cluster (7D/I). By 12.25dpc, these TuJ1-expressing neuronal fibers have extended from the progenitor cluster towards the distal tip of the genital tubercle accompanied by *Sox10-H2BVenus+* cells in close proximity to the extending neuronal fibers (Figure 7D/I). By 12.5dpc, the neuronal processes extending past the developing pelvic ganglia clusters have grown longer and more prominent (Figure 7E/J). This analysis confirms the presence of sizable aggregates of HuC/D positive progenitors bilaterally alongside the genital tubercle that are apparent when viewed from a lateral perspective (Figure 7E).

**Figure 7.**
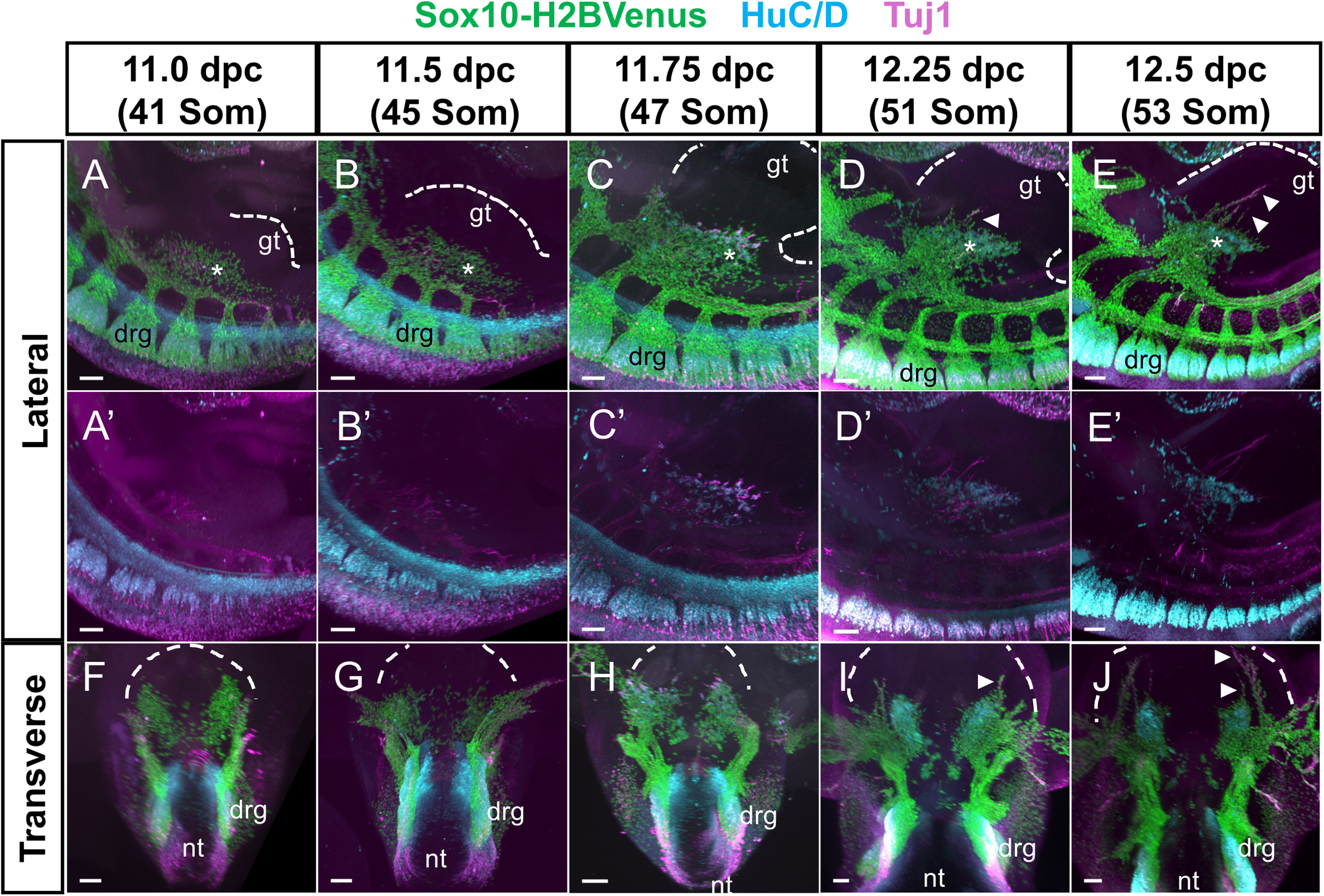
LSFM reveals precise spatiotemporal assembly of pelvic ganglia precursor aggregates and initiation of neurogenesis. LSFM imaging of *Sox10-H2BVenus* embryos at representative timepoints between 11.0dpc and 12.5dpc reveals the spatial dynamics of the initial colonization of the genital tubercle by neural crest-derived progenitors. (A-E) lateral perspective of the sacral region of embryos at 11.0dpc, 11.5dpc, 11.75dpc, 12.25dpc, and 12.5dpc. *Sox10*-expressing migratory neural crest progenitor cells are shown in green, immunostaining for HuC/D is shown in cyan, and immunostaining for TuJ1 is shown in magenta. Asterisks (*) signify the population of neural crest-derived cells on the genital tubercle (dashed lines) which will give rise to the pelvic ganglia. (F-J) Transverse perspectives of the images shown in (A-E) show the bilateral colonization of the genital tubercle (dashed lines) by neural crest-derived cells. By 12.25dpc, streams of neural crest-derived progenitor cells have moved towards the distal aspect of the genital tubercle accompanied by neural fibers (D,E,I,J, white arrowheads). Abbreviations: **drg,** dorsal root ganglia; **nt,** neural tube. Scale bars = 200um.

## DISCUSSION

In this study, we produced a three-dimensional light sheet fluorescence microscopy atlas of pelvic ganglia formation in the mouse, ranging initial stages when sacral neural crest progenitors first colonize the urogenital sinus to mature, functional ganglia morphology in young adult mice. We adapted the CUBIC tissue clearing method for use in mouse LUT tissue, allowing for optimal visualization of pelvic innervation when applied to tissue from *Sox10-cre; R26R-tdTomato* and *Sox10-H2BVenus* mice. Using immunohistochemistry for markers of neuronal identity, we were further able to visualize the spatial organization of neuronal somata and processes. In total, our imaging approach provided novel three-dimensional perspective and insight for the complexity of pelvic innervation in the mouse.

In LSFM imaging of LUTs from male and female mice at P28, we observed that the distribution of neuronal somata in the pelvic ganglia is far more extensive than has been previously appreciated. While the field has described the mouse MPG as a centralized body of neurons, we find that there are many HuC/D-expressing neurons outside of this central cluster, present anteriorly along the pelvic and hypogastric nerves, posteriorly down the length of the cavernous nerves, and ventrally on the surface of the urethra. In our imaging, we also observed that the bladder-proximal accessory ganglia are multi-lobular and there is a sizeable cluster of neurons present along the accessory fibers, which we have termed the ‘intermediate ganglia’. As has been noted previously,^16^ we find that the MPG diverges from bladder-proximal ganglia populations postnatally. Whether these intermediate and accessory ganglia serve a unique or focused purpose in the mature postnatal LUT is yet to be understood. Our spatial mapping of these populations acts as an important step in parsing out the functional significance of subpopulations within the mouse pelvic ganglia and sets the stage for defining the factors that regulate this separation of distinct pelvic ganglia domains.

Using our established tissue clearing and LSFM protocols, we also produced three-dimensional renderings of pelvic ganglia morphology at key embryonic timepoints. By imaging pelvic ganglia in late gestation, we find that neuronal distribution around the LUT is initially far less centralized than in the mature pelvic ganglia structure. At 15.5dpc, we observed several neuronal somata present on the ventral surface of the urethra, and these cells increase in number by 18.5dpc. However, at postnatal timepoints, the number of neuronal cells on the ventral urethra decreases as the pelvic ganglia are consolidated into major, accessory, and intermediate populations. We hypothesize that this transient distribution of neuronal cell bodies around the LUT may play a functional role in tissue remodeling and morphogenesis as reproductive organs mature. In the male, prostate formation occurs between 15.5dpc and 18.5dpc and is accompanied by ingrowth of tyrosine hydroxylase expressing nerve fibers,^21^ and so neuronal somata on the ventral urethra may be necessary for proper development of this organ.

Lastly, we investigated the migratory entry routes of sacral neural crest-derived progenitor cells which colonize the developing LUT. As in previous investigations,^3^ we observed that neural crest progenitors begin to populate the lateral sides of the genital tubercle swelling by 11.0dpc. We identify the time between 11.5dpc and 11.75dpc as a key stage for initiation of neurogenesis among sacral neural crest progenitors that will form the pelvic ganglia. Our studies suggest that there is a regionality to the distribution of HuC/D+ cells within the aggregate of sacral pelvic ganglia progenitors with greater density of HuC/D+ cells in the ventral aspect. Further molecular analysis of this short developmental window will be highly informative for defining regulatory cues that control and guide neurogenesis in the pelvic ganglia. Future studies of transcriptional dynamics between neural crest-derived progenitors and surrounding tissue during this period of migration and neurogenesis to uncover the complex nature of how and why progenitors organize in this way to form the pelvic ganglia will be highly informative for clinical efforts seeking to innervate artificial bladder scaffolds.^22,23^ Interestingly, our studies reveal that the aggregate of sacral neural crest-derived progenitors that form the pelvic ganglia are quite distinct and spatially resolved from the sympathetic chain that is located more dorsal.

In all, the work presented here will act as a necessary framework for further understanding pelvic ganglia formation, as the spatial resolution we have achieved in our efforts has been missing in prior studies of pelvic innervation. For decades, our knowledge of pelvic ganglia morphology and organization has been largely limited to what was learned from morphological studies of larger mammals or based on removal of only the major pelvic ganglia population away from the LUT. ^4,5,24,25,26,27,28^ These prior studies have been instrumental in building our understanding of neuronal cell types and the diversity of neuropeptide signaling in the MPG population. However, added spatial perspective is necessary to strengthen our understanding of how various cell types contribute to the many functions of the LUT. Our LSFM atlas offers an improved appreciation for the complexity of pelvic innervation, which in turn will be significant for subsequent studies of pelvic ganglia origin, diversity, and function.

## LIMITATIONS

In our selection and optimization of tissue clearing methods, we applied CUBIC clearing to mouse LUT tissue. We chose to apply CUBIC, which uses aqueous solutions, due to the limited amount of tissue distortion we observed in our samples and constraints in using organic solvents in our local imaging system. Since the pelvic ganglia is adjacent to, but not embedded within, the LUT organs, we took measures to ensure that tissue clearing would not disrupt the relative spatial organization of innervating bodies and innervated tissue. One major limitation of this study is that CUBIC clearing does not achieve the same optical clarity as alternative organic solvent-based methods. Therefore, we were unable to attain image resolution through the entire volume of our samples, especially in samples from older mice with more complex tissue. We worked around this obstacle because the pelvic ganglia are largely superficial populations of cells and we were able to dissect away extraneous tissue (e.g. fat, prostate) from the LUT, guided by our endogenous fluorescent reporters.

## METHODS

### Mouse husbandry

All experimental protocols were approved by the Institutional Animal Care and Use Committee (IACUC) at Vanderbilt University Medical Center. All mice were housed in a modified barrier facility on a 14-hour on, 10-hour off-light cycle in high-density caging (Lab Products Inc., #10025) with water ad libitum. Weanlings and non-breeders received standard diet (LabDiet #5L0D), and breeders received high-energy diet (LabDiet #5LJ5). Tg ^(Sox^^10-cre)1Sout^ (MGI:5897484), hereafter referred to as *Sox10*-cre, were maintained as a congenic line on C3HeB/FeJ (C3Fe, Jackson Stock# 00658) by breeding heterozygous transgenic males with wildtype C3Fe females.^15^ *Sox10*-cre females were crossed with B6.Cg-*Gt(ROSA)26Sor^tm14(CAG-tdTomato)Hze^*/J, hereafter *R26R-tdTomato*, mice to visualize neural crest derivatives of pelvic ganglion based on *tdTomato* expression. Tg ^(*Sox10*-HIST2H2BE/Venus)ASout^ mice (MGI: 3769269), hereafter called *Sox10-H2BVenus*, were maintained by crosses with female C3Fe.^20^

*Sox10-cre; R26R-tdTomato* and *Sox10-H2BVenus* mice were fluorescently screened for *tdTomato* and *Venus* expression, respectively. Tissue samples were collected from animals expressing either of the fluorescent proteins, and automated genotyping was performed by Transnetyx. *Sox10-cre* and *Sox10-H2BVenus* transgene sequences were confirmed using primers amplifying the Sp6 and T7 vector promoters and internal BAC-specific sequences. Primer sequences are listed in Table 1.

**Table 1:**
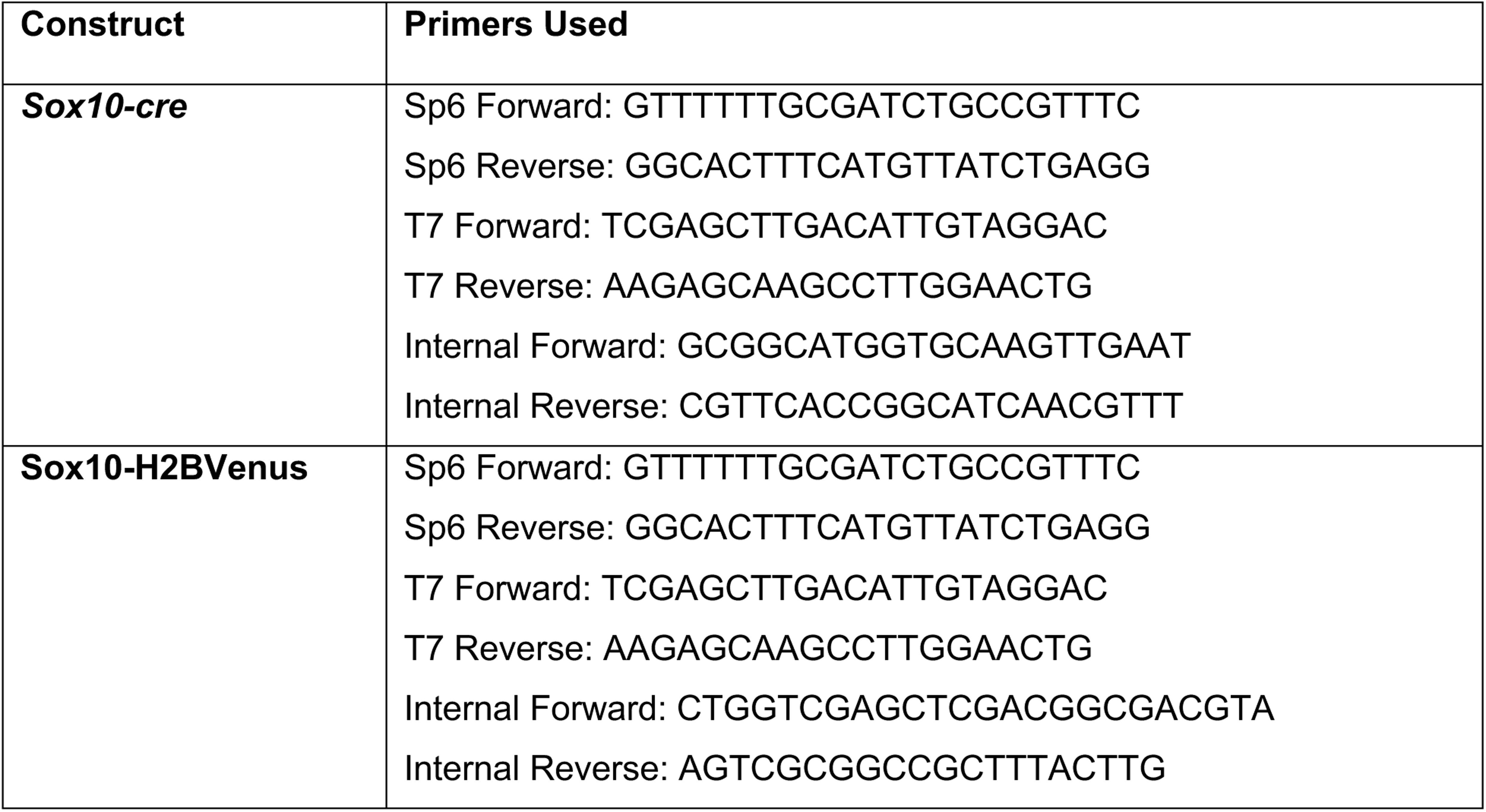
Primer sequences used for genotyping.

### Tissue collection and processing

LUT tissue was collected from postnatal *Sox10-cre; R26R-tdTomato* animals at P2, P10, and P28. Prior to dissection, anesthetized animals were subjected to transcardiac perfusion with fixative (4% paraformaldehyde in 1X PBS) using a Variable-Flow Peristaltic pump (Fisher). P2 animals were perfused with 10 mL of fixative, P10 animals were perfused with 20 mL of fixative, and P28 animals were perfused with 40 mL of fixative. Post-perfusion, the LUT was removed from the body, and fluorescence-guided dissection was performed to remove any extraneous tissue present on top of the *tdTomato*-positive pelvic ganglia populations. Following dissection, additional drop tissue fixation was performed in 4% paraformaldehyde overnight at 4°C. Tissue was then washed out of fix with 1x PBS and stored in 0.1% Sodium Azide in 1x PBS.

*Sox10-cre; R26R-tdTomato* embryos were collected at 12.5, 15.5 and 18.5dpc. For this effort, timed matings were performed between *Sox10-cre* females and *R26R-tdTomato* males according to established methods, with the morning of copulatory plug observation designated as 0.5dpc. Upon embryo collection from pregnant dams, developmental stage was confirmed by comparing forelimb and hindlimb morphology to known standards. Embryos were then bisected at the level of the liver, and extraneous tissue was removed to expose the developing pelvic ganglia. Embryonic tissue was then fixed overnight in 4% paraformaldehyde with 6% sucrose and 0.5% Triton X-100 in 1x PBS. Tissue was then washed out of fix with 1x PBS and stored in 0.1% Sodium Azide in 1x PBS.

*Sox10-H2BVenus* embryos were collected at 11.0, 11.5. 11.75, 12.25, and 12.5dpc using a similar method to above. Timed matings were performed between C3Fe females and *Sox10-H2bVenus* males. Embryo collection was performed at various times between the morning of 11dpc and the evening of 12dpc. Embryos were staged by comparing somite counts and gross morphology to known standards.^29^ Forelimbs were taken for genotyping and embryos were otherwise kept intact. This tissue was fixed overnight in 4% paraformaldehyde with 6% sucrose and 0.5% Triton X-100 in 1x PBS. Tissue was then washed out of fix with 1x PBS and stored in 0.1% Sodium Azide in 1x PBS.

Replicate counts for *Sox10-cre; R26RtdTomato* and *Sox10-H2BVenus* samples are outlined in Supplementary Tables 1 and 2. All replicates listed supported the morphological observations outlined in the text.

### Tissue clearing and Immunohistochemical labeling

Prior to LSFM imaging, tissue was processed in wholemount through CUBIC clearing and immunohistochemistry. The standard CUBIC processing pipeline includes two aqueous solutions: CUBIC R1 is used for tissue delipidation, and CUBIC R2 for refractive index matching. The CUBIC R1 solution consists of 25% Urea, 25% N,N,N′,N′-tetrakis(2-hydroxypropyl)ethylenediamine (Quadrol), and 15% Triton X-100 by weight in deionized water. The CUBIC R2 solution consists of 50% Sucrose, 25% Urea, and 0.1% Triton X-100 by weight in deionized water.

All clearing steps were performed at 30°C, as we have observed that higher temperatures quench *tdTomato* fluorescence. Tissue was first incubated in a 1:1 dilution of CUBIC R1 in deionized water for 24 hours. Following this incubation, tissue was transferred into undiluted CUBIC R1 for 5-7 days, with solution changes daily. Once tissue appeared optically clear, samples were washed out of CUBIC R1 with 1x PBS + 0.6% Triton X-100 and processed through immunohistochemistry.

Before immuno-staining, samples were incubated at 4°C overnight in blocking solution containing 5% Normal Donkey Serum, 1% Bovine Serum Albumin, and 0.5% Saponin in 1x PBS. Primary and secondary antibodies were diluted as listed in Tables 2 and 3 in 5% Normal Donkey Serum, 1% Bovine Serum Albumin, and 0.6% Triton X-100 in 1x PBS. Tissue was incubated with primary antibodies for 24 hours at 4°C, followed by 4 washes out of primary antibody for 60 minutes each with 1x PBS + 0.6% Tx-100. Samples were then incubated in secondary antibody for 6 hours at 4°C. To remove unbound secondary antibody, samples were washed 10 times for at least 30 minutes each in 1x PBS + 0.6% Tx-100, with at least one of these washes being overnight. Samples were then post-fixed for 1 hour in 4% PFA. Samples were then washed out of post-fix three times for at least 10 minutes each in 1x PBS + 0.6% Tx-100. Finally, samples were incubated for 20 minutes in autofluorescence quenching solution consisting of 0.5 mM Cupric Sulfate in 50 mM Ammonium Acetate.

**Table 2:** Primary antibodies used for analysis.

| Target antigen | Host | RRID | Supplier | Dilution | Incubation |
|---|---|---|---|---|---|
| Hu C/D | Human | AB_2813895 | Gift from Vanda Lennon | 1:10,000 | 24 hours at 4ºC |
| Class III beta-tubulin, neuronal (TuJ1) | Rabbit (polyclonal) | AB_291637 | Covance Cat# PRB-435P-100 | 1:5,000 | 24 hours at 4ºC |

**Table 3:** Secondary antibodies for detection of primary antibodies.

| **Target antigen** | **Host** | **RRID** | **Supplier** | **Dilution** | **Incubation** |
|---|---|---|---|---|---|
| Anti-human, Alexa Fluor 647 | Donkey | AB_2340578 | Jackson ImmunoResearch Inc., 709-605-149 | 1:1000 in *Sox10-cre; R26R-tdTomato* samples;<br>1:500 in *Sox10-H2BVenus* samples | 6 hours at 4ºC |
| Anti-rabbit,<br>Cy3 | Donkey | AB_2307443 | Jackson ImmunoResearch Inc., 711-165-152 | 1:750 | 6 hours at 4ºC |

Following immuno-staining, samples were incubated in CUBIC R2 solution at 30°C for 7-14 days, with solution changes daily. Samples regained opacity while in immuno-staining solutions, and therefore they were kept in CUBIC R2 until they were optically clear. Once clear, tissue was mounted for LSFM.

### Mounting for Light Sheet Imaging

Since embryonic and lower urinary tract tissue lack rigidity, even after fixation, samples were mounted in agarose to keep them in place during LSFM. For this approach, a 1.5% solution of low-melt agarose in Milli-Q water was prepared. A 3cc syringe was trimmed at the base of the barrel near the tip and filled with approximated 2 mL of warm 1.5% agarose via aspiration. The syringe was then inverted so that it was resting on the plunger handle with the trimmed open end facing up. Samples were then transferred into a basin containing the warmed agarose solution for equilibration and to remove any air bubbles. Following this rinse in agarose, samples were positioned inside the 3cc syringe and held in place using laboratory forceps while the agarose solidified. Once the agarose was solid, the sample was released. The agarose-mounted sample was then removed by depressing the plunger of the syringe, and the ends of the agarose were trimmed to be just longer than the height of the sample. The sample was then incubated for 2-3 additional days in CUBIC R2 at 30°C clear the agarose.

Just before LSFM imaging, samples were super-glued onto a small, flat sheet of metal. Small magnets were also glued inside the end of a glass capillary which was compatible with the sample holder of the microscope. This allowed for the agarose-embedded samples to be mounted inside the microscope using magnets attached to a short wooden dowel.

### Light Sheet Imaging and Image Processing

Light sheet fluorescence microscopy imaging in this study was performed on a Zeiss Z.1 microscope equipped with a 2X pco.edge 4.2 CLHS camera with a liquid cooled sCMOS sensor. Images were acquired using a 5x NA 0.16 objective lens. Samples were imaged while submerged in CUBIC R2 solution (refractive index = 1.47) inside the imaging chamber. Acquisition parameters were set using the Zeiss ZEN Black imaging software. All wholemount LSFM images were collected as tiled z-stacks using the following laser wavelengths: 488, 561, 647nm. Data from each laser wavelength was collected as separate tracks, switching tracks after each z-stack. Tile overlap was set to 10%, with a 2 second delay between z-stack collection. Laser power and exposure time were adjusted depending on thickness, angle, and clarity of each sample, and therefore these settings were not maintained across samples. The Pivot Scan function was utilized to reduce shadows when passing the light sheet through samples. Zoom settings were maintained at 1x for all images.

Following image acquisition, file outputs from the Zeiss ZEN software (.czi) were converted into Imaris files (.ims) using the Imaris FileConverter 10.2.0 software. The tiled LSFM images were then aligned and stitched using the Imaris Stitcher 10.2.0 software. Finally, image analysis and figure production was performed in Imaris 10.2.0 using the ‘Snapshot’ and ‘Animation’ functions.

## ACKNOWLEDGEMENTS

The authors thank the staff of the Cell Imaging Shared Resource (CISR) Core at Vanderbilt for advice and assistance in light sheet imaging. HuC/D antibody was a kind gift from Vanda Lennon (Mayo Clinic).

## Data Availability

Original image files and 3-D visualizations (segmentations) will be available for viewing or download on sparc.science.

