## Supplemental Materials for "Organ scale Three-dimensional atlas of developing pelvic innervation for the mouse lower urinary tract"

**Supplementary Table 1: Summary of LSFM imaging replicates supporting findings in *Sox10-cre; R26R-tdTomato* samples.**

| Timepoint | Sex | # Replicates supporting conclusions |
|---|---|---|
| 12.5dpc | M/F | 6 |
| 15.5dpc | M | 3 |
| 15.5dpc | F | 4 |
| 18.5dpc | M | 3 |
| 18.5dpc | F | 3 |
| P2 | M | 2 |
| P2 | F | 3 |
| P10 | M | 4 |
| P10 | F | 3 |
| P28 | M | 3 |
| P28 | F | 3 |

**Supplementary Table 2: Summary of LSFM imaging replicates supporting findings in *Sox10-H2BVenus* samples.**

| Timepoint | # Replicates supporting conclusions |
|---|---|
| 11.0dpc | 4 |
| 11.5dpc | 3 |
| 11.75dpc | 4 |
| 12.25dpc | 3 |
| 12.5dpc | 3 |

**Supplementary Figure S1. Non-neural crest-derived glandular tissue expresses the *Sox10-cre* transgene postnatally.** Wholemount LSFM imaging of *Sox10-cre; R26R-tdTomato* male lower urinary tracts reveals robust expression of the *tdTomato* reporter (magenta) in the glandular urethra (A) and in distal portions of prostatic lobes (A'). Abbreviations: b, bladder; u, urethra; mpg, major pelvic ganglia. Scale bars = 800um (A), 150um (A').

**Supplementary Figure S2. The female accessory ganglia are also comprised of distinct ganglia lobules.** Wholemount LSFM imaging of the lower urinary tract from a female *Sox10-cre; R26R-tdTomato* mouse at P28 shows complexity of bladder-proximal ganglia organization. (A). Neural crest lineages (magenta) are shown relative to neuronal somata labeled by immunostaining for HuC/D (cyan). To improve visualization, spatial distribution of the accessory ganglia was defined according to HuC/D fluorescent signal (A'), and HuC/D positive neuronal somata outside of the defined region were masked out of the image to isolate the accessory ganglia (A''). Three-dimensional observation of the accessory ganglia both from lateral (A'') and ventral (B) perspectives shows discrete ganglia populations present at the bladder neck (dashed lines). Larger populations are marked with asterisks (*) and smaller populations are marked with apostrophes ('). Abbreviations: **AG,** accessory ganglia; **IG,** intermediate ganglia; **b,** bladder; **A,** Anterior; **P,** posterior; **D,** dorsal; **V,** ventral; **L,** lateral; **M,** medial. Scale bars = 100um.

**Supplementary Figure S3. Spatially distinct pelvic ganglia populations also separate postnatally in female mice.** Wholemount lower urinary tracts from *Sox10-cre; Rosa*$^{Tom}$ female mice imaged via LSFM at postnatal timepoints show a continuum of change in pelvic ganglia organization after birth. Neural crest lineages (magenta) are shown relative to neuronal somata labeled by immunostaining for HuC/D (cyan). At P2 (A), there is a small separation between the major pelvic ganglia and the accessory ganglia at the bladder neck (populations marked by asterisk, separated by dotted line). By P10 (B), there is a larger space between the centralized major pelvic ganglia and the bladder-proximal accessory ganglia (populations marked by asterisks, separated by dotted line). Finally, at P28 (C), mature morphology of pelvic innervation is established, and a diffuse network of nerve fascicles connect separate populations of major pelvic ganglia, intermediate ganglia, accessory ganglia (populations marked by asterisks, separated by dotted lines). Abbreviations: **b,** bladder; **u,** urethra. Scale bars = 300um.

**Supplementary Figure S4. Neuronal populations expand dorsally and ventrally during late gestation in female embryos.** Wholemount lower urinary tracts from *Sox10-cre; Rosa$^{Tom}$* male embryos at 15.5 dpc (A) and 18.5 dpc (B) show changes in neuronal cell body localization in the pelvic ganglia which occur at late embryonic timepoints. Neural crest lineages (magenta) are shown relative to neuronal somata labeled by immunostaining for HuC/D (cyan). At 15.5 dpc, a centralized pelvic ganglia structure has formed (A'). By 18.5 dpc, just before birth, pelvic ganglia neurons have expanded to the ventral and dorsal region of the urethra. (B', white arrowheads). Scale bars = 500um (A,B), 200um (inset zooms).

Supp Figure 1_Mehaffey et al., 2026

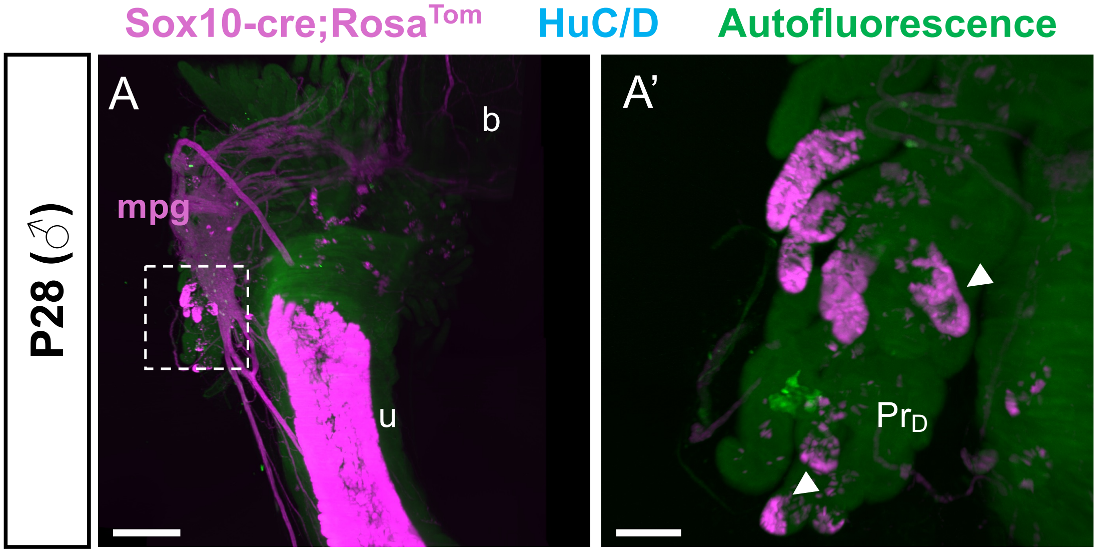

### Supp Figure 2_Mehaffey et al., 2026

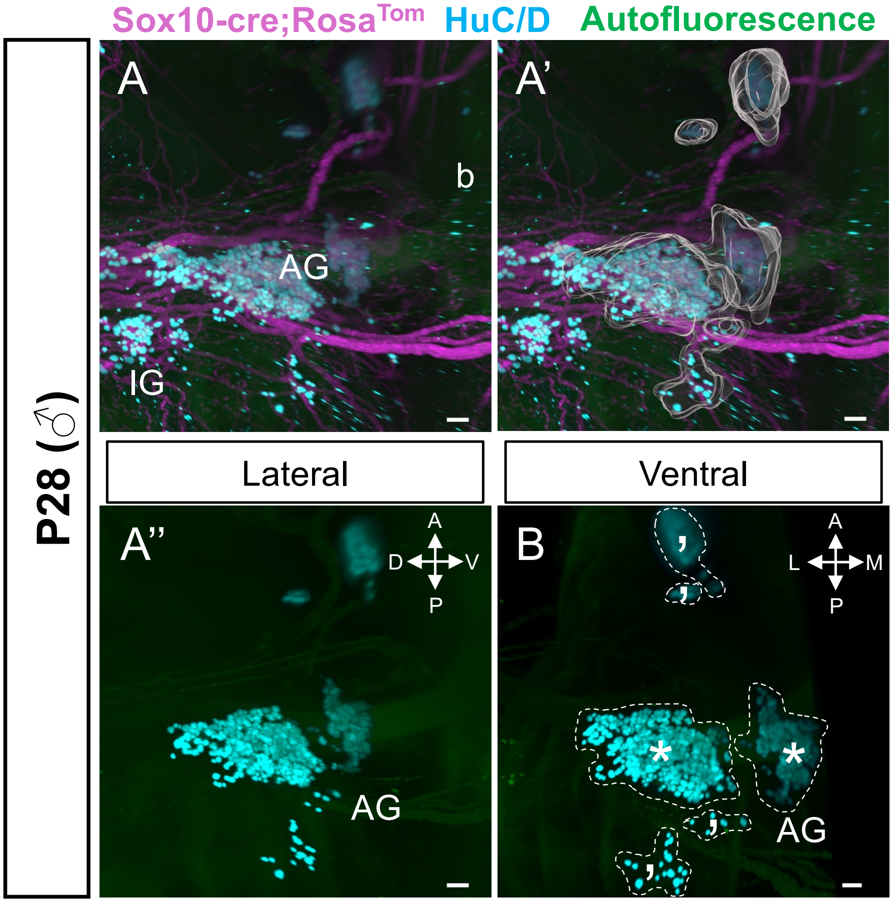

Supp Figure 3_Mehaffey et al., 2026

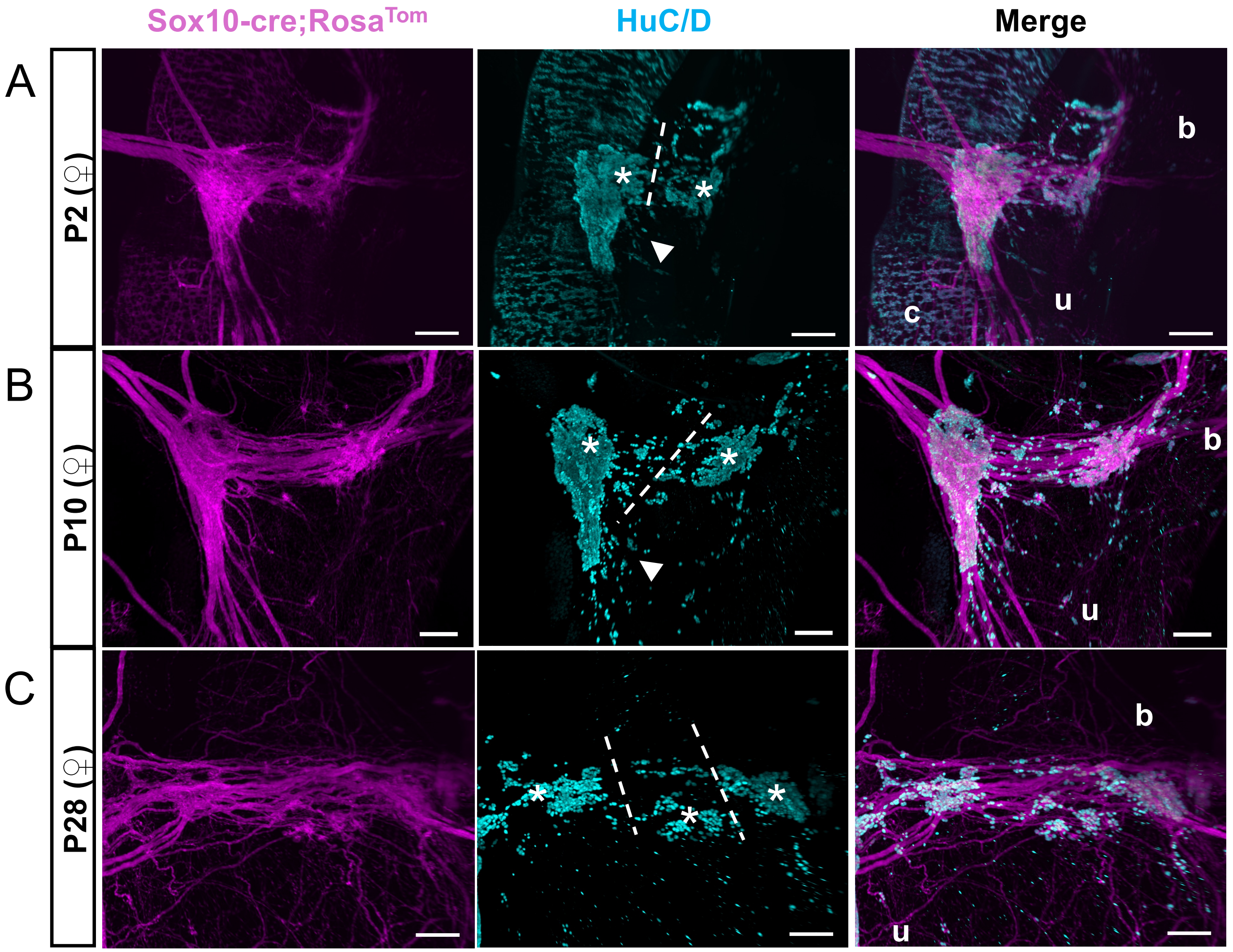

Supp Figure 4_Mehaffey et al., 2026

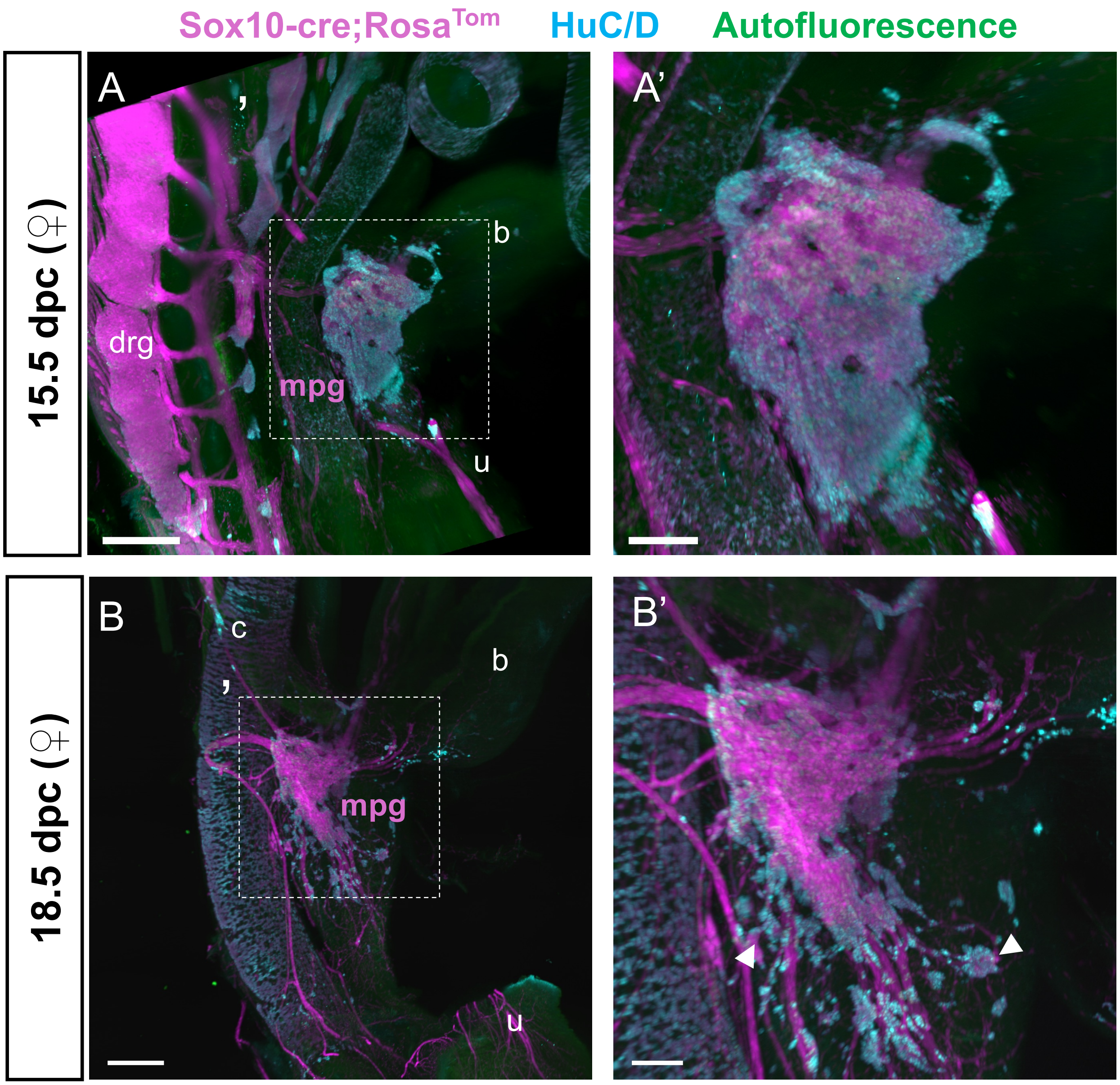
